# SC-MEB: spatial clustering with hidden Markov random field using empirical Bayes

**DOI:** 10.1101/2021.06.05.447181

**Authors:** Yi Yang, Xingjie Shi, Wei Liu, Qiuzhong Zhou, Mai Chan Lau, Jeffrey Chun Tatt Lim, Lei Sun, Joe Yeong, Jin Liu

## Abstract

Spatial transcriptomics has been emerging as a powerful technique for resolving gene expression profiles while retaining tissue spatial information. These spatially resolved transcriptomics make it feasible to examine the complex multicellular systems of different microenvironments. To answer scientific questions with spatial transcriptomics and expand our understanding of how cell types and states are regulated by microenvironment, the first step is to identify cell clusters by integrating the available spatial information. Here, we introduce SC-MEB, an empirical Bayes approach for spatial clustering analysis using a hidden Markov random field. We have also derived an efficient expectation-maximization algorithm based on an iterative conditional mode for SC-MEB. In contrast to BayesSpace, a recently developed method, SC-MEB is not only computationally efficient and scalable to large sample sizes but is also capable of choosing the smoothness parameter and the number of clusters. We performed comprehensive simulation studies to demonstrate the superiority of SC-MEB over some existing methods. We applied SC-MEB to analyze the spatial transcriptome of human dorsolateral prefrontal cortex tissues and mouse hypothalamic preoptic region. Our analysis results showed that SC-MEB can achieve a similar or better clustering performance to BayesSpace, which uses the true number of clusters and a fixed smoothness parameter. Moreover, SC-MEB is scalable to large ‘sample sizes’. We then employed SC-MEB to analyze a colon dataset from a patient with colorectal cancer (CRC) and COVID-19, and further performed differential expression analysis to identify signature genes related to the clustering results. The heatmap of identified signature genes showed that the clusters identified using SC-MEB were more separable than those obtained with BayesSpace. Using pathway analysis, we identified three immune-related clusters, and in a further comparison, found the mean expression of COVID-19 signature genes was greater in immune than non-immune regions of colon tissue. SC-MEB provides a valuable computational tool for investigating the structural organizations of tissues from spatial transcriptomic data.

## 1 Introduction

Recent advances in spatial transcriptomics (ST) have allowed researchers to simultaneously measure transcriptome-wide gene expression at near single-cell resolution while the spatial information for each measurement is retained [Burgess, 2019]. These spatially resolved transcriptomics have deepened our understanding of how cell types and states are regulated by tissues microenvironment, e.g., of the human brain [Maynard et al., 2021], mouse brain [Shah et al., 2016, Alon et al., 2021], and mouse embryo [Lohoff et al., 2020], among others. The technologies used for resolving spatial gene expression can largely be classified as either imaging-based or next-generation-sequencing-based methods [Waylen et al., 2020]. Imaging-based methods, which were developed to study spatial complexity, are based on fluorescent in situ hybridization (FISH) and include smFISH [Lyubimova et al., 2013], seqFISH [Eng et al., 2017], and MERFISH [Xia et al., 2019]. Although FISH-based methods are capable of capturing both RNA quantity and position, they are limited by their throughput scalability and accuracy in measuring gene expression levels. However, multiple next-generation-sequencing-based methods have been developed to facilitate high-throughput analysis, including Geo-seq [Chen et al., 2017], Slide-seq [Rodriques et al., 2019] and, more recently, the commercial 10x Genomics Visium system [Ståhl et al., 2016]. Emerging ST technologies offer new opportunities to investigate the spatial patterns of gene expression for many applications, such as cell type identification, tissue exploration, and differential expression analysis. Among these applications, cell-type clustering is the first problem that needs to be addressed.

Similar to single-cell RNA-seq data, ST data contains excessive amounts of zeros or “drop-outs” [Qiu, 2020]. Recently, many academics have argued that drop-outs are mostly due to biological variation, such as cell-type heterogeneity, rather than technical shortcomings [Svensson, 2020]. Kim et al. Kim et al. [2020] suggested that clustering analysis should be performed before imputing or normalizing the data. To overcome the curse of dimensionality due to high-throughput spatial gene expression, clustering is often preceded by standard dimension reduction procedures, e.g., principal component analysis (PCA), *t*-distributed stochastic neighbor embedding [Van der Maaten and Hinton, 2008], and uniform manifold approximation and projection [McInnes et al., 2018].

In ST datasets, the majority of existing clustering methods, e.g., k-means [Kriegel et al., 2017] and Gaussian mixture models (GMM) [Bishop, 2006], do not consider the available spatial information. To allow additional spatial information to be incorporated into ST datasets, several methods have been recently developed, including the hidden Markov random field model implemented in the *Giotto* package [Zhu et al., 2018, Dries et al., 2019] and a fully Bayesian model with a Markov random field (MRF), BayesSpace [Zhao et al., 2021]. Given the spatial coordinates for each transcriptome-profiled spot, spatial clustering methods achieve better classification accuracy. For example, Zhao et al. [2021] showed that BayesSpace improved the resolution and achieved better classification accuracy for manually annotated human brain samples. However, these methods have certain limitations. First, BayesSpace is a fully Bayesian method based on Markov chain Monte Carlo; therefore, it is not computationally scalable for ST data with high resolution. Second, smoothness is an essential parameter of MRF-based methods and largely determines the proximity of the neighboring spots [Tolpekin and Stein, 2009]. BayesSpace takes this smoothness parameter as fixed and, thus, cannot choose the optimal value that best fits a given dataset. Third, without optimizing the smoothness parameter, one cannot apply any model selection methods to obtain the optimal number of clusters. In practice, the number of clusters in ST datasets is usually unknown before the follow-up analysis, and the preferred method would be to automatically choose the number of clusters.

To address these limitations, we propose a method of <u>S</u>patial <u>C</u>lustering using the hidden <u>M</u>arkov random field based on <u>E</u>mpirical <u>B</u>ayes (SC-MEB) to model a low-dimensional representation of a gene expression matrix that incorporates the spatial coordinates for each measurement. In contrast to existing methods [Dries et al., 2019, Zhao et al., 2021], SC-MEB is not only computationally efficient and scalable to larger sample sizes but also accommodates adjustments to the smoothness parameter and the number of clusters. We derived an efficient expectation-maximization (EM) algorithm based on an iterative conditional mode (ICM) and further selected the number of clusters for SC-MEB based on the modified Bayesian information criterion (MBIC) [Wang et al., 2009]. We demonstrated the effectiveness of SC-MEB over existing methods through comprehensive simulation studies. We then applied SC-MEB to the clustering analysis of three ST datasets. Using a 10x Genomics Visium dataset from human dorsolateral prefrontal cortex tissues that were manually annotated, we showed that the performance of SC-MEB was comparable or better than that of BayesSpace, even though the latter uses the “true” number of clusters and a prespecified, fine-tuned smoothness parameter. Using a large MERFISH dataset from mouse hypothalamic preoptic region, we demonstrated the better clustering performance as well as the scalability of SC-MEB. We further applied SC-MEB and alternative methods to analyze ST data of a colon tissue from a patient with colorectal cancer (CRC) and COVID-19. We performed follow-up differential expressions analysis using the clustering results from SC-MEB and BayesSpace, and the heatmap of identified signature genes showed that SC-MEB clustering results were more reasonable and interpretable. Using pathway analysis, we identified three immune-related clusters, and the mean expression of COVID-19 signature genes were further compared between immune and non-immune regions of a colon sample.

## 2 Materials and Methods

### 2.1 Problem Formulation

The SC-MEB consists of three major stages (Fig. 1A). First, PCA is conducted on the log-transformed expression of the highly variable genes to obtain the top principal components (PCs) (Fig. 1B). Next, spatial clustering is performed using PCs for each spot. Finally, downstream analyses, such as differential expression analysis, can be performed to obtain signature genes for each cluster (Fig. 1D).

**Figure 1:**
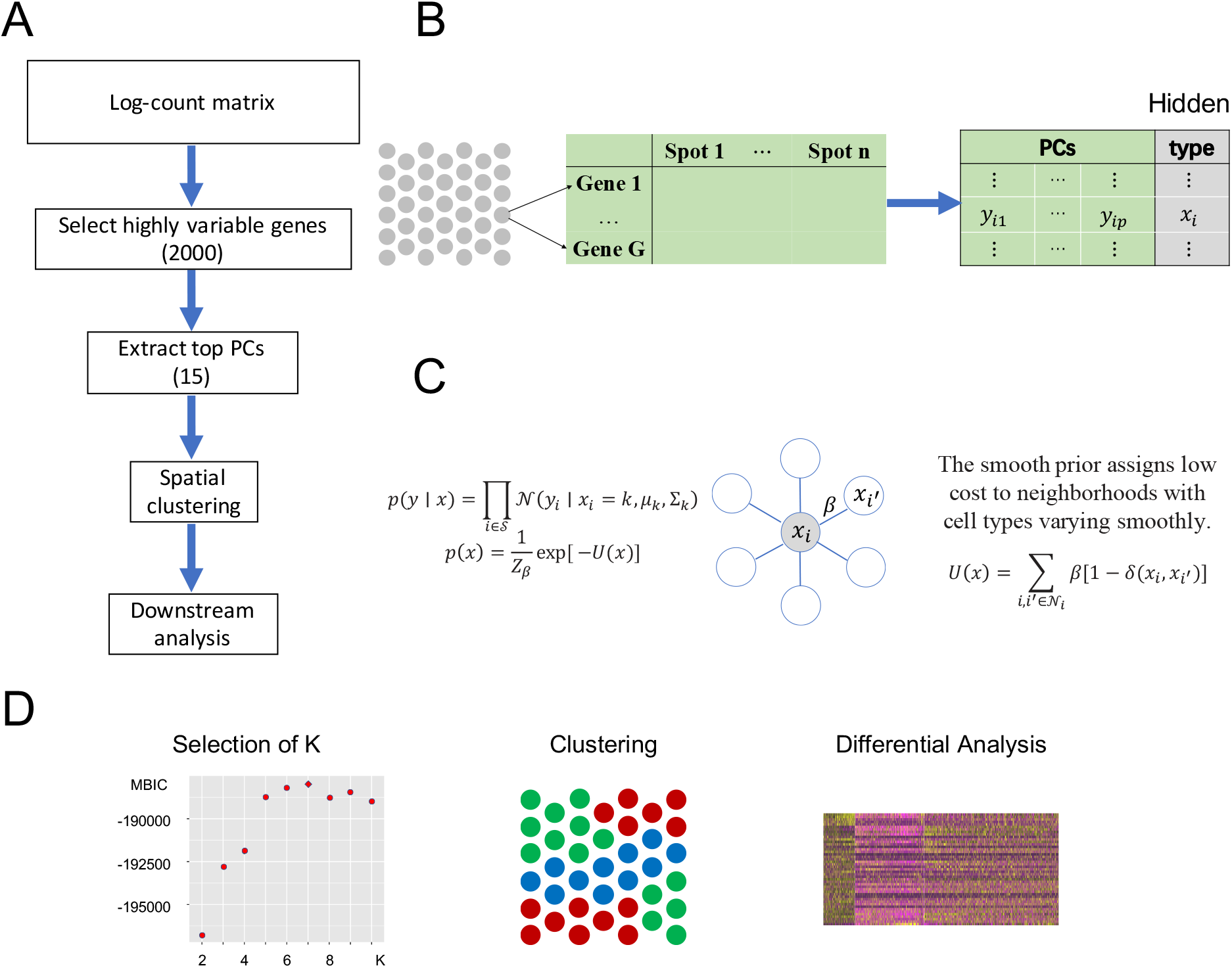
SC-MEB workflow. **A**. The SC-MEB workflow mainly comprises the following steps, data preprocessing, spatial clustering using the hidden MRF model, a series of down-stream analyses. **B**. Data preprocessing: log-transformation, dimension reduction. **C**. The hidden MRF model. For the Visium dataset, we usedd six neighborhoods for each spot. **D**. The SC-MEB outputs: a scatter plot of MBIC for all K, a tissue plot with spots colored by clustering, a heatmap of DEGs.

Our spatial clustering method builds on a two-level hierarchical probabilistic model (Fig. 1C). Briefly, for spot *i*, the first level specifies the conditional probability of the low-dimensional representation (e.g., top PCs) of its gene expression *y*_*i*_ given an unknown label *x*_*i*_ ∈ {1,…,*K*}, where *K* is the number of clusters. In SC-MEB, we assume that given the labels for each spot, a *d*-dimensional representation *y*_*i*_ is mutually independent among all spots, and its distribution within a given cluster *k* can be written as

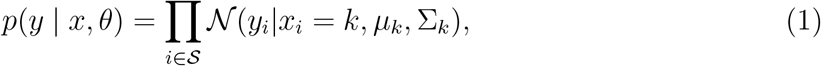

where *θ* = {*μ*_*k*_, Σ_*k*_ : *k* = 1,…, *K*}, and *μ*_*k*_ and Σ_*k*_ denote the mean and covariance matrix for cluster *k*, respectively.

The second level of SC-MEB depicts the prior probability of the hidden labels, and an MRF prior is implemented to encourage smoothness among spots. In other words, spots of the same cluster can be in close proximity. As spots in Visium are primarily arranged on hexagonal lattices, the neighborhood of each spot is defined by applying a proximity threshold. To promote smoothness within spot neighborhoods, we use the Potts model [Potts, 1952] for the hidden labels. The Potts model is well known as a statistical model for use with complex systems with nearest neighbor interactions. Essentially, it views the total energies *U*(*x*) as a summation of pairwise interaction energies with neighbors, where a positive parameter *β* represents the strength of interactions. Specifically, the Potts model promotes spatial smoothness by penalizing cases in which neighboring spots are assigned to different cluster labels. The hidden random field *x* is assumed to be

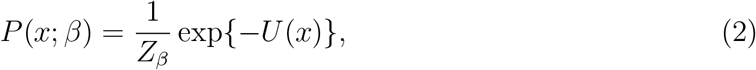

where 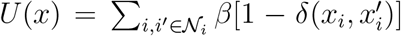, *δ* is the delta function, and *Z*_*β*_ is a normalization constant that does not have a closed form. When all labels on a neighborhood take the same value, meaning that the hidden *x* is locally smooth, it incurs no neighborhood cost; otherwise, if they are not all the same, a positive cost is incurred, and the amount of cost is controlled by parameter *β*. Thus, parameter *β* controls the smoothness in latent labels; the larger the *β*, the spatially smoother the latent labels. When *β* is zero, SC-MEB reverts to the method that does not consider spatial information, i.e., GMM. Combining two levels of SC-MEB, (1) and (2), we denote *ϕ* = (*θ, β*) the parameter space.

As the smoothing parameter *β* does not have an explicit updated form, SC-MEB adaptively selects *β* via a grid search strategy. That is, the SC-MEB model is trained with a prefixed *β* using an efficient ICM-EM scheme [Cuadra et al., 2005] that incorporates a pseudo-likelihood maximization step, as in the ICM method of Besag [1974]. The optimal *β* is the value that maximizes the marginal log-likelihood. In a similar way, the marginal log-likelihood can be evaluated for a sequence of *K*. Then, MBIC [Wang et al., 2009] is applied to choose the optimal number of clusters in a data-driven manner (Fig. 1D). Please refer to Supplementary for more details about the MBIC used in this case.

### 2.2 ICM-EM Algorithm

The parameter is estimated through an iterative-conditional-mode-based expectation-maximization (ICM-EM) algorithm [Cuadra et al., 2005]. Here, we assume *K* is known.

In the ICM step, the estimate of *x* is obtained by maximizing its posterior with respect to *x*_*i*_ coordinately:

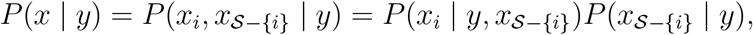

where *i* = 1,…, *n*, until converge [Besag, 1986]. Given initial values of *x, ϕ*, and observed *y*, we have the updated equation:

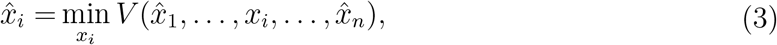

where

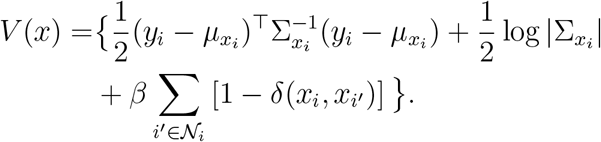

In the expectation (E) step, instead of using the original complete likelihood, which is difficult to evaluate, the following pseudo-likelihood is used:

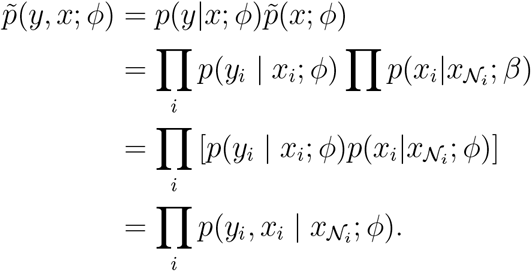

With the optimal conditional distribution of *x* (details in Supplementary), we have

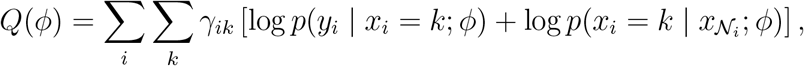

where *γ*_*ik*_ is the responsibility that component *k* has for explaining the observation *y*_*i*_, which is defined as follows:

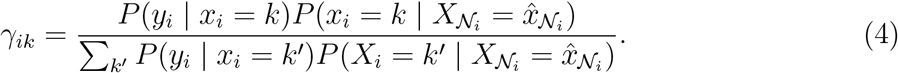

By taking partial derivatives of *Q*(*ϕ*) with respect to the parameter *θ* and setting them to zero, we obtain the updated equations in the maximization (M) step:

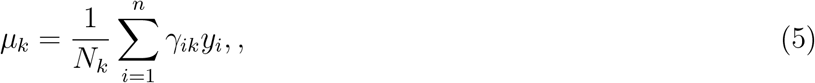

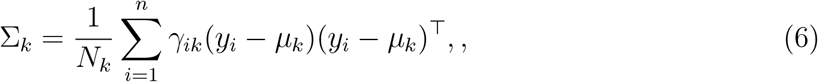

where 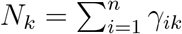. Since there is no closed-form solution for *β*, we optimize the smoothness parameter *β* via a grid search strategy:

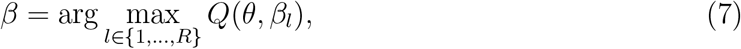

where the sequence (*β*_1_,…, *β*_*R*_) is a vector of 20 evenly spaced points in the interval [0, 4].

The ICM-EM algorithm iterates the ICM step and M step until convergence. Further details on the ICM-EM algorithm are provided in the Supplementary Material.

### 2.3 Methods for comparison

We conducted comprehensive simulations and real data analysis to gauge the performance of different methods for clustering a low-dimensional representation of a gene expression matrix, including both non-spatial and spatial clustering methods.

In detail, we considered the following non-spatial clustering methods: (i) k-means implemented in the R package *stats*, available at CRAN; (ii) GMM implemented in the R package *mclust*, available at CRAN; (iii) Louvain implemented in the R package *igraph*, available at https://igraph.org/r/. In addition, we compared the clustering performance of spatial methods: (i) SC-MEB implemented in the R package *SC*.*MEB*, available at https://github.com/Shufeyangyi2015310117/SC.MEB; (ii) BayesSpace implemented in the R package *BayesSpace*, available at Bioconductor; (iii) HMRF implemented in the *Giotto* package, available at http://spatialgiotto.rc.fas.harvard.edu/.

### 2.4 Preprocessing of ST datasets

The Visium spatial transcriptomics [Zhao et al., 2021] data were aligned and quantified using Space Ranger downloaded from 10x Genomics official website against the GRCh38 human reference genome also from 10x Genomics official website. For all datasets, we applied log-transformation of the raw count matrix using library size [Lun et al., 2016, McCarthy et al., 2017]. Then, we performed PCA on the 2,000 most highly variable genes. In the clustering analysis, we chose the top 15 PCs from the study datasets as the input for SC-MEB as well as for the alternative methods.

### 2.5 ST datasets

#### 2.5.1 Human dorsolateral prefrontal cortex (DLPFC)

Maynard et al Maynard et al. [2021] used recently released ST technology, the 10x Genomics Visium platform, to generate spatial maps of gene expression matrices for the six-layered DLPFC of the adult human brain that are provided in the *spatialLIBD* package. They also provided manual annotations of the layers based on the cytoarchitecture. In their study, they profiled the spatial transcriptomics of human postmortem DLPFC tissue sections from 12 samples, with a median depth of 291 M reads for each sample, corresponding to a mean 3,462 unique molecular indices and a mean 1,734 genes per spot.

#### 2.5.2 Mouse hypothalamic preoptic region (MHPR)

Moffitt et al. Moffitt et al. [2018] used the combination of MERFISH with scRNA-seq to profile the gene expression of 1 million cells in situ that reveals neuronal populations in the preoptic region of 36 mouses, each with distinct molecular signatures and spatial organizations. Specifically, the MHPR dataset contains expression values of 161 genes in 1,027,848 cells. To demonstrate the scalability, we perform joint clustering for all cells of the 36 samples. The spatial locations for each sample are offset so that cells of different samples are not neighbors. Here we add 10,000 to row and column coordinates to achieve this. Sample 1 further contains 6 slices, we refer them as Sample 1-1 to Sample 1-6. We further analyze all cells in Sample 1 as well as each of these six slices. The number of cells for each dataset are summarized in Table 4. On average, there are six neighbors for every cells determined by their Euclidean distance orders.

#### 2.5.3 Human colon tissue adjacent to colorectal cancer (CRC)

The colon tissue was from a 45-year-old South Asian male who was diagnosed with COVID-19 on April 16, 2020. As previously described [Cheung et al., 2021], the patient had experienced mild upper respiratory tract symptoms throughout the course of the disease. He was confirmed COVID-19-negative after two consecutive nasopharyngeal swabs on and May 9 and 10, 2020, and was discharged from the isolation facility on May 10, 2020. During hospital admission, further investigation involving computed tomography scanning and colonoscopy revealed the presence of a large circumferential malignant mass in the cecum. Histology of the biopsies confirmed that the patient had invasive colorectal cancer stage II T3N0. He underwent laparoscopic right hemicolectomy on May 18, 2020, 9 days after testing negative for COVID-19. He recovered uneventfully and was discharged on May 21, 2020. Using this sample, we profiled the spatial transcriptomics using the 10x Genomics Visium platform. In summary, it has a depth of 143 million reads for a total of 2988 spots within the tissue and a median 492 genes per spot.

### 2.6 Evaluation metrics

We evaluated the clustering performance by adjusted Rand index (ARI) [Schütze et al., 2008]. The general formula for ARI is as follows

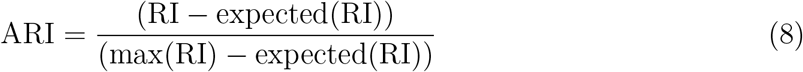

where RI is the Rand index [Rand, 1971], and max (RI) and expected (RI) are the maximum value and the expected value of RI, respectively. Assuming that *n* is the number of spots in an ST dataset. *U* = {*u*_1_,…, *u*_*i*_,…, *u*_*R*_}∈ ℝ^*n*^ and *V* = {*v*_1_,…, *v*_*j*_,…, *v*_*C*_}∈ ℝ^*n*^ represent two clustering labels for *n* spots, where *R* and *C* are the corresponding numbers of clusters in *U* and *V*, respectively. Denoting *n*_*ij*_ as the number of spots belonging to both classes *u*_*i*_ and *v*_*j*_, and *n*_*i·*_ and *n*_*·j*_ as the number of spots in classes *u*_*i*_ and *v*_*j*_, respectively; then ARI (8) is defined as

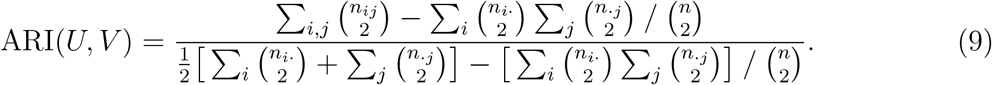

As the expected value of RI for two random partitions does not take a constant value and is concentrated within a small interval, ARI is a corrected version of RI to avoid these drawbacks [Hubert and Arabie, 1985]. Note that ARI lies between -1 and 1 and takes a value of 1 when the two partitions are equal up to a permutation. Obviously, a larger ARI value indicates a higher similarity between two partitions. In the simulation, ARI was used to measure the similarity between the estimated partition and the true one. In the analysis of the DLPFC dataset [Maynard et al., 2021], manual annotations based on additional experiments and computational results were available. ARI was used to measure the similarity between labels from the estimated partition and the manually annotated clusters.

## 3 Results

### 3.1 Simulation settings

Using simulations, we compared the clustering performance of SC-MEB and with five other methods, including k-means, GMM, Louvain, BayesSpace, and Giotto. For *k*-means, BayesSpace, and Giotto, we considered the true number of clusters *K*, and its two nearest numbers, *K* − 1 and *K* + 1, as the number of clusters had to be manually specified for these two methods. For all other methods, the number of clusters was selected automatically. The smoothness parameter *β* of BayesSpace was fixed at 3 by default, while *β* of SC-MEB was optimized with a grid search. We compared the clustering performances using ARI for all methods, in which we ran 50 replicates in each setting.

In Example I, the labels for spots were randomly generated. In detail, for a 70 *×* 70 squared lattice with 4,900 spatial spots, we generated cluster labels for each spot from the *K*-states Potts model (as shown in Eqn. (2)) with *β* ∈ [1, 1.3] using the R package *GiRaF*. The number of neighbors was set to be 4, and the number of true clusters *K* was set to 3, 5, or 7. We then considered two distributions for low-dimensional PCs: a mixture of Gaussian and a mixture of Student’s-*t* distributions. The number of PCs was set to either 10 or 15. The mean *μ*_*k*_ and the covariance matrix Σ_*k*_ for each component *k* are listed in Supplementary Tables S1-S4.

In Example II, labels for spots were obtained from real data analysis. In detail, we used the inferred cluster labels from SC-MEB (*K* = 8) of colon data as the true labels for all 2,988 spots. PCs were randomly generated in the same way as in Example I. The mean *μ*_*k*_ and the covariance matrix Σ_*k*_ for each component *k* are provided in Supplementary Tables S5-S6.

In the above examples, all *K* components had different covariance matrices. Because BayesSpace adopts a strategy in which all components have a shared covariance, we further conducted additional simulations with equal covariance matrices.

### 3.2 Performance of SC-MEB in comparison with other methods in simulation studies

In Example I, when PCs were from a mixture of Gaussian distributions, SC-MEB was more powerful than all other methods (Fig. 2A). BayesSpace had a smaller ARI, i.e. poorer concordance between predicted and true clustering assignment, than SC-MEB, even when the true number of clusters was used as input. The inferior performance of BayesSpace was due to its lack of adaptation to the smoothness parameter *β*. The other methods, Giotto, GMM, *k*-means, and Louvain, achieved lower ARIs. When PCs were from a mixture of Student’s *t*-distributions, assumptions of BayesSpace were satisfied. As shown in Fig. 2B, using BayesSpace with the correct number of clusters showed the best performance. Even though SC-MEB was miss-specified in this setting, it still achieved a high ARI that was larger than that of BayesSpace with miss-specificed *K* and other methods. Note that most components in the mixture of Student’s *t*-distributions simulated here are *t*(5) and *t*(6), which are reasonably close to Gaussian. This demonstrates the robust performance of SC-MEB when there is moderate miss-specification of distributions. We note that if the data are far from the Gaussian component, the performance of SC-MEB will degenerate. The results from other settings (Supplementary Fig. S1-S2) prompted similar conclusions.

**Figure 2:**
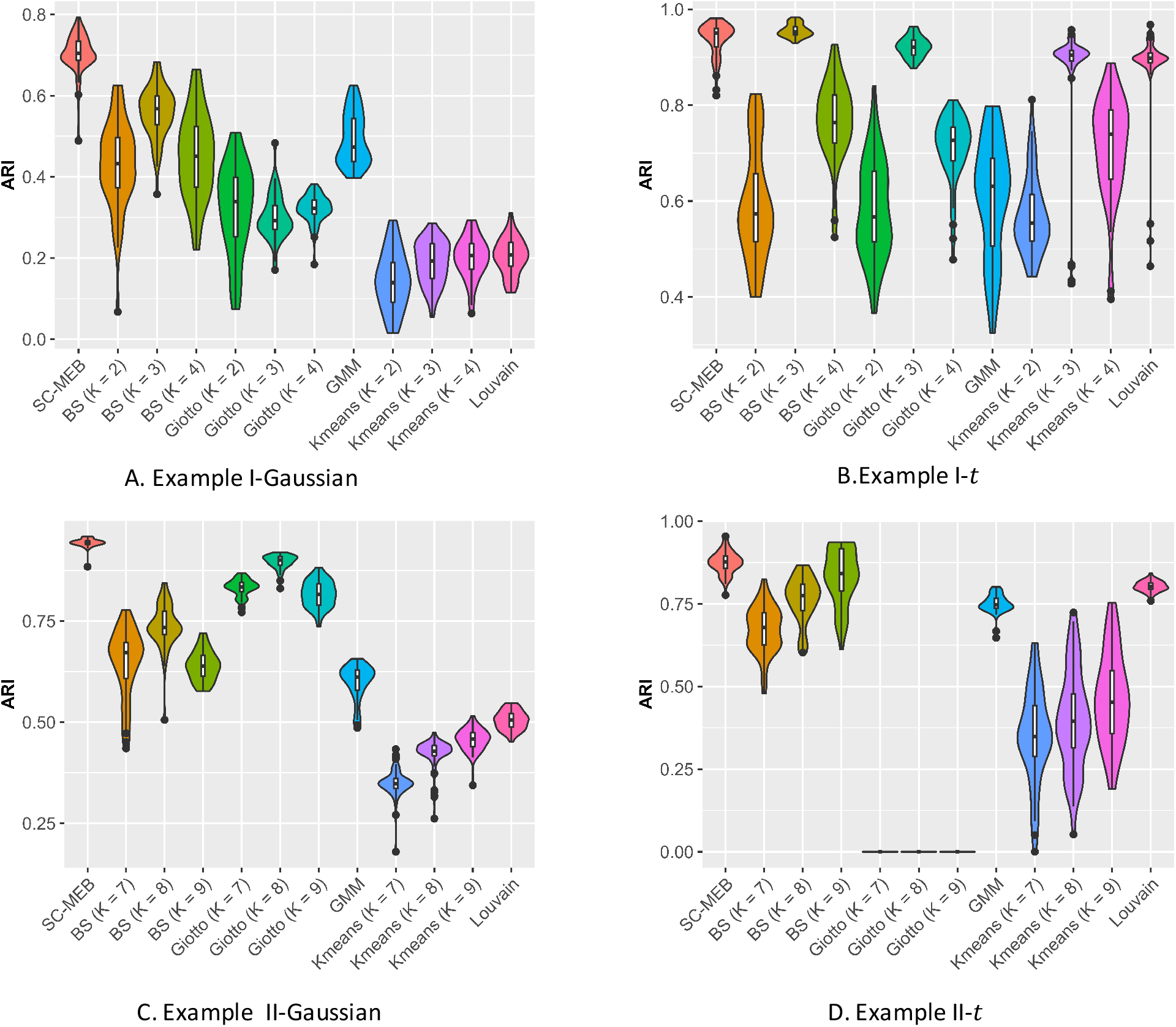
Summary of clustering accuracy of the six methods in the analysis of simulated data. **A**. Example 1, Gaussian: PCs were sampled from a Gaussian mixture model. **B**. Example 1, t: PCs were sampled from a Student’s-*t* mixture model. **C**. Example 2, Gaussian: PCs were sampled from a Gaussian mixture model. **D**. Example 2, t: PCs were sampled from a Student’s-*t* mixture model.

In Example II, the comparative results (Fig. 2C and 2D) were largely consistent with the results obtained in Example I. Specifically, SC-MEB was more powerful than all the other methods. The performance of BayesSpace was the next most powerful, and *k*-means had the worst performance. The results obtained from other settings (Supplementary Fig. S5A-S5B, S6A-S6B) led to similar conclusions.

Finally, we considered the above two examples under BayesSpace’s assumption that all *K* components share a common covariance. The results are shown in (Supplementary Fig. S3-S4, S5C-D, and S6C-D). As is shown, the ARI of SC-MEB was comparable with that of BayesSpace, and both demonstrated better performance than the other methods.

All simulations were conducted on a computer with a 2.1 GHz Intel Xeon Gold 6230 CPU and 16 GB memory. SC-MEB was computationally more efficient than BayesSpace. In all simulations, SC-MEB toke approximately 8 minutes to complete the analysis for 10 combinatorial values in *K*, while BayesSpace required about 25 minutes for prefixed *K* and *β* and up to 600 times more computation time than SC-MEB for fixed combinatorial values of *K* and *β*. To better demonstrate the computational efficiency and scalability of SC-MEB, we conducted additional simulations with an increasing sample size *n*. In Fig. 3, we can see that the computation time of SC-MEB for a fixed number of iterations increased almost linearly with increasing sample size, taking about 0.5 hours to run 50 iterations for a dataset with 200K spots. Thus, SC-MEB can be used to perform clustering analysis for ST datasets with a higher resolution than other methods.

**Figure 3:**
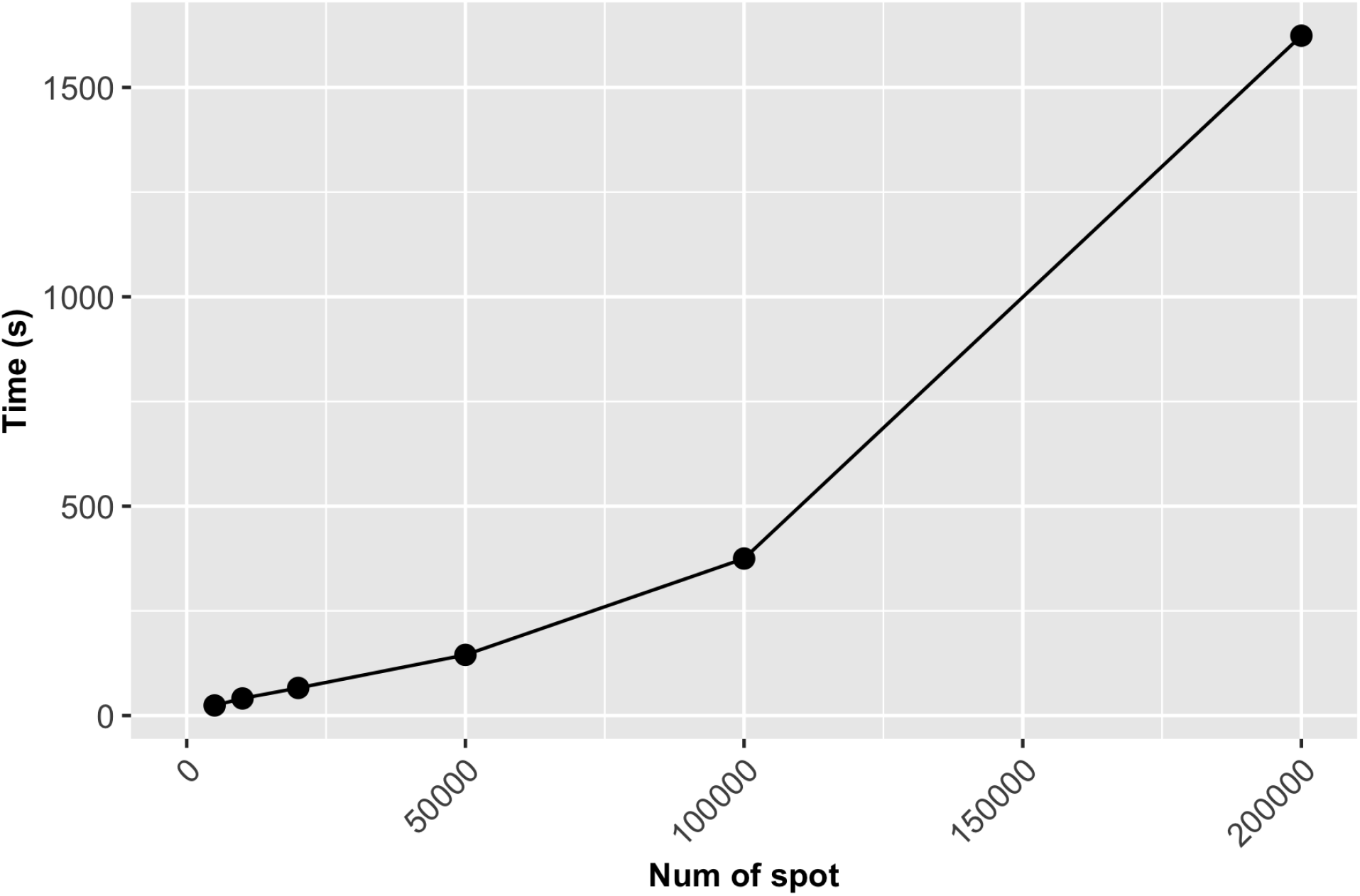
The computation time of SC-MEB increases linearly with sample size. The number of iterations was set to 50 for the different sample sizes.

### 3.3 Benchmark clustering performance with real datasets

To evaluate the clustering performance of SC-MEB with real datasets, it was applied to the DLPFC and MHPR datasets and its clustering performance was compared with that of alternative methods. Specifically, we first obtained the top 15 PCs from the 2000 most highly variable genes in DLPFC dataset and all 161 genes in MHPR dataset, respectively. Then we performed clustering analysis with all methods, except k-means. As BayesSpace and Giotto cannot choose the number of clusters *K*, the K was set to the number of clusters in the manual annotations. All other methods selected the number of clusters automatically.

Table 1 shows the ARI values for 12 DLPFC samples, where the manually annotated layers were taken as the “ground truth”. SC-MEB clearly outperformed BayesSpace in the analysis of six samples, and vice versa for the other five samples. In the analysis, BayesSpace took both the “true” number of clusters (from the manual annotations) and the prefixed fine-tuned *β* as input. In this case, the proposed SC-MEB achieved the similar performance without the prior information. Additionally, SC-MEB achieved the best clustering performance among the methods that can select (*K*) automatically. Table 2 compares the computational times required for all methods. The speed of SC-MEB was almost 200 times faster than that of BayesSpace, and comparable to GMM and Giotto. Louvain was the fastest method, but its clustering accuracy was inferior.

**Table 1:** Clustering accuracy for DLPFC dataset. ARI values were evaluated by comparing manual annotations against cluster labels from SC-MEB and alternative methods for all 12 samples.

| ID | SC-MEB | BayesSpace | GMM | Giotto | Louvain |
| --- | --- | --- | --- | --- | --- |
| 151507 | **0.42** | 0.33 | 0.40 | 0.33 | 0.32 |
| 151508 | **0.44** | 0.36 | 0.33 | 0.34 | 0.25 |
| 151509 | **0.52** | 0.44 | 0.29 | 0.35 | 0.30 |
| 151510 | 0.39 | **0.43** | 0.31 | 0.33 | 0.28 |
| 151669 | 0.32 | **0.41** | 0.22 | 0.25 | 0.20 |
| 151670 | **0.43** | **0.43** | 0.19 | 0.21 | 0.26 |
| 151571 | **0.42** | 0.38 | 0.23 | 0.40 | 0.36 |
| 151672 | 0.44 | **0.77** | 0.14 | 0.38 | 0.27 |
| 151673 | 0.49 | **0.55** | 0.29 | 0.37 | 0.29 |
| 151674 | **0.43** | 0.33 | 0.29 | 0.29 | 0.33 |
| 151675 | 0.31 | **0.41** | 0.24 | 0.32 | 0.24 |
| 151676 | **0.39** | 0.32 | 0.26 | 0.26 | 0.25 |

**Table 2:** Comparison of computation times (s) of five methods in the analysis of DLPFC. Note, that for SC-MEB, we used a K sequence from 2 to 10 and a sequence of *β* from 0 to 4.

| ID | SC-MEB | BayesSpace | GMM | Giotto | Louvain |
| --- | --- | --- | --- | --- | --- |
| 151507 | 23.00 | 7015.95 | 46.16 | 33.10 | 2.86 |
| 151508 | 23.55 | 5471.57 | 36.26 | 20.48 | 6.97 |
| 151509 | 23.56 | 4660.43 | 53.22 | 27.05 | 6.00 |
| 151510 | 26.62 | 4288.56 | 39.50 | 24.27 | 7.57 |
| 151669 | 23.79 | 5840.10 | 20.96 | 20.81 | 3.49 |
| 151670 | 18.96 | 4917.43 | 37.94 | 16.35 | 5.26 |
| 151571 | 24.54 | 3889.07 | 31.01 | 20.28 | 8.82 |
| 151672 | 27.91 | 3793.58 | 22.44 | 21.92 | 5.49 |
| 151673 | 18.92 | 5984.89 | 40.20 | 35.30 | 1.49 |
| 151674 | 19.80 | 5344.68 | 36.82 | 23.57 | 3.34 |
| 151675 | 18.93 | 3863.19 | 27.90 | 26.01 | 4.22 |
| 151676 | 18.93 | 3609.71 | 37.44 | 28.07 | 3.24 |

The ARI and computation times of the five methods for MHPR dataset are provided in Table 3 and 4. Obviously, SC-MEB has the best clustering performance. The ARI of BayesSpace for each dataset is lower than SC-MEB. And for large datasets such as all cells and cells in Sample 1, it cannot work. The Giotto and Louvain can work well for each dataset, but their ARI is smaller than SC-MEB.

**Table 3:** Clustering accuracy for MHPR dataset. ARI values were evaluated by comparing molecularly annotations against cluster labels from SC-MEB and alternative methods for all 8 samples.

| ID | SC-MEB | BayesSpace | GMM | Giotto | Louvain |
| --- | --- | --- | --- | --- | --- |
| All | **0.47** | - | 0.25 | 0.19 | 0.36 |
| 1 | **0.51** | - | 0.20 | 0.21 | 0.37 |
| 1-1 | 0.44 | 0.24 | 0.23 | 0.22 | **0.46** |
| 1-2 | **0.46** | 0.41 | 0.20 | 0.25 | 0.41 |
| 1-3 | **0.49** | 0.31 | 0.20 | 0.26 | 0.41 |
| 1-4 | **0.52** | 0.35 | 0.22 | 0.23 | 0.42 |
| 1-5 | 0.39 | 0.31 | 0.22 | 0.24 | **0.40** |
| 1-6 | **0.49** | 0.31 | 0.21 | 0.25 | 0.41 |

**Table 4:** Comparison of computation times (s) of five methods in the analysis of MHPR. Note that for SC-MEB, we used a sequence of *K* from 8 to 27 and a sequence of *β* from 0 to 4.

| ID | $n$ | SC-MEB | BayesSpace | GMM | Giotto | Louvain |
| --- | --- | --- | --- | --- | --- | --- |
| All | 1027848 | 16647 | - | 107914 | 11940 | 45884 |
| 1 | 73665 | 1390 | - | 14288 | 744 | 287 |
| 1-1 | 11738 | 318 | 16152 | 2192 | 153 | 10 |
| 1-2 | 12672 | 349 | 17238 | 2267 | 172 | 13 |
| 1-3 | 12556 | 357 | 19071 | 2332 | 170 | 17 |
| 1-4 | 11763 | 326 | 15459 | 2293 | 150 | 10 |
| 1-5 | 12306 | 383 | 16818 | 2130 | 160 | 10 |
| 1-6 | 12620 | 332 | 18366 | 2234 | 164 | 14 |

### 3.4 Spatial clustering of in-house CRC sample

To apply SC-MEB in the analysis of an in-house colon sample from a patient suffeed from CRC and COVID-19, we first obtained the top 15 PCs, as described for the DLPFC dataset. The spatial clustering performed by SC-MEB was compared with that of other methods. Because SC-MEB, Giotto, and Louvain selected eight clusters as the optimal number (*K*), we also ran BayesSpace with eight clusters. The computational times for SC-MEB, BayesSpace, GMM, Giotto, and Louvain were 24.66, 5324.48, 46.26, 49.88, and 0.69 seconds, respectively.

The clustering results obtained using the different methods are shown in Fig. 4. In general, the pattern of clustering assigned by SC-MEB was similar to that of GMM, but the latter retained more noisy spots. In addition, the results from SC-MEB and BayesSpace had stronger spatial patterns than those of the other methods.

**Figure 4:**
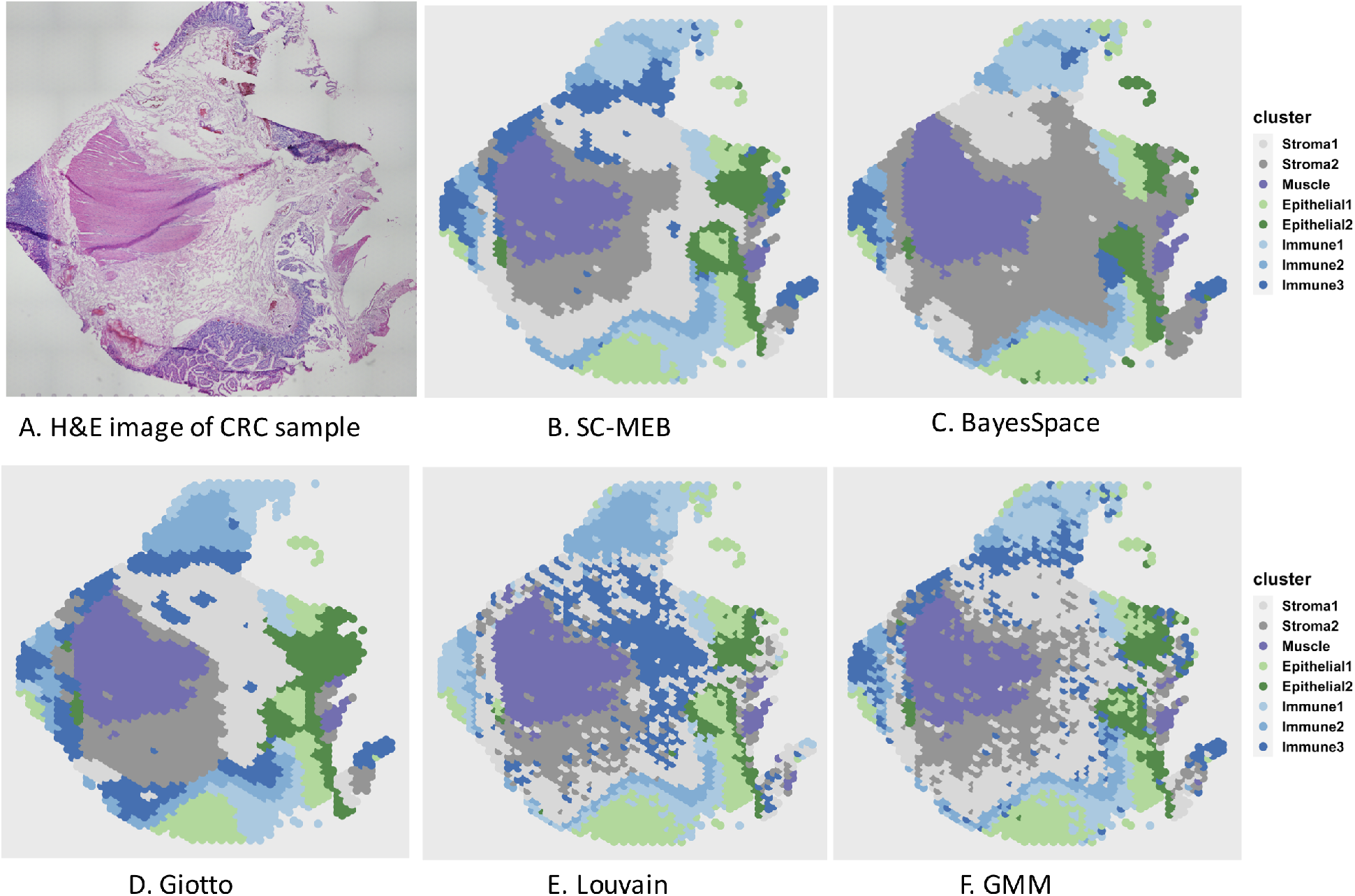
Clustering results for a colon sample. (A) Original H&E-stained tissue image for the colon sample. (B-F) Heatmaps for clustering assignments in the colon sample using the proposed SC-MEB, BayesSpace, Giotto, Louvain, and GMM, respectively. The eight clusters identified included two stromal regions, a muscle region, two epithelial-cell regions, and three immune-cell regions.

By checking PanglaoDB [Franzén et al., 2019] for signature genes identified via *differential expression (DE) analysis* and with the help of the H&E staining shown in Fig. 4A, we were able to identify regions of muscle, stroma, epithelial, and immune cells. As shown in Fig. 4B-F, all methods except BayesSpace returned good partitions for the muscle region, which were visually verified with the H&E staining (Fig. 4A). The epithelial regions identified by BayesSpace were much smaller than those identified by SC-MEB, in which a large proportion of the epithelial regions in Fig. 4B-C were classified as stromal regions (stroma 2) by BayesSpace. The immune regions identified by BayesSpace were larger at the 9 and 12 o’clock positions but smaller at the 6 o’clock position in Fig. 4C than those identified by SC-MEB (Fig. 4B) and GMM (Fig. 4F). Strikingly, a large proportion of the regions identified as stroma 1 by both SC-MEB and GMM were classified as stroma 2 by BayesSpace. Even though stroma 1 and stroma 2 are both stromal regions, one can observe clear differences in the normalized expression of signature genes for these two clusters (Fig. 5). These observations illustrate the possible over-smoothing behavior of BayesSpace, while SC-MEB was able to recover the fine structure of tissues.

**Figure 5:**
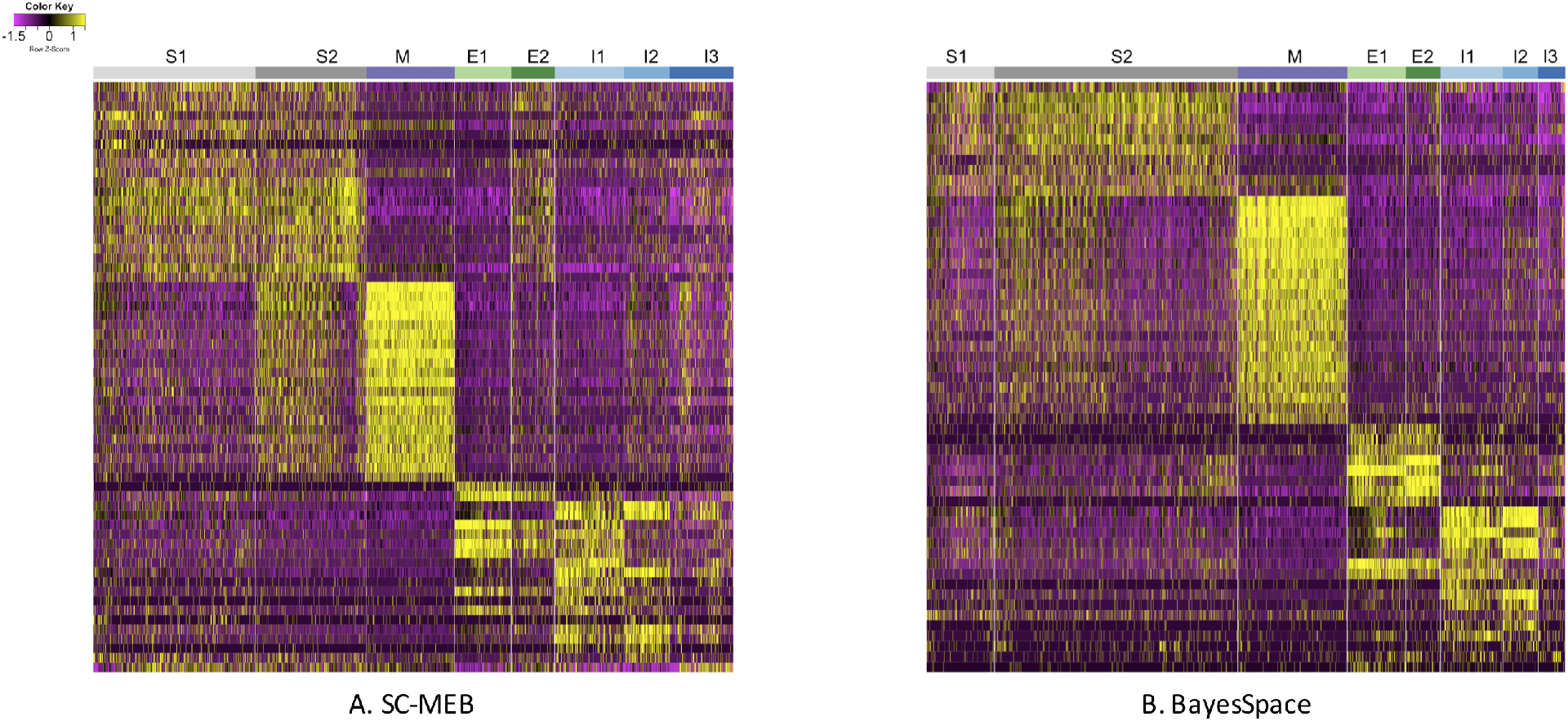
Heatmaps of normalized expression of signature genes identified in the differential expression analysis based on two clustering analysis methods: (A) SC-MEB and (B) BayesSpace. In both subfigures, S1 and S2 represent Stroma 1 and 2, respectively; M is Muscle; E1 and E2 are Epithelial 1 and 2, respectively; and I1, I2, and I3 are Immune 1, 2, and 3, respectively.

#### 3.4.1 Differential expression analysis of the identified clusters

As true labels for all spots were not available for the colon dataset, we could not quantitatively evaluate the clustering performance. For the clustering results of SC-MEB and BayesSpace, we further performed DE analysis comparing an identified cluster with all others using the *BPSC* package [Vu et al., 2016] for log-normalized expression. Using the partition results from SC-MEB, we identified 180, 158, 128, 94, 145, 95, 137, and 84 genes that were differentially expressed for stroma 1 and 2; muscle; epithelial 1 and 2; and immune 1, 2, and 3, respectively, with a false discovery rate of *<* 0.05. The details of all differentially expressed genes identified by SC-MEB and BayesSpace are provided in Supplementary Tables S7 and S8. We further restricted the number of signature genes by choosing those with log-fold changes larger than 0.5. Finally, we obtained a total of 62 and 57 signature genes for SC-MEB and BayesSpace, respectively.

Fig. 5 shows the heatmap of normalized expression for the signature genes identified in the DE analysis by SC-MEB (Fig. 5A) and BayesSpace (Fig. 5B), respectively. Clearly, with BayesSpace, the normalized expression of signature genes in stroma 2 could be further divided into two sub-clusters, and the expression pattern in the second sub-cluster was very similar to that of stroma 1. This misclassification is also apparent when comparing Fig. 4B and Fig. 4C, as a large proportion of the regions identified as stroma 1 by SC-MEB were identified as stroma 2 by BayesSpace. The findings obtained using SC-MEB demonstrated that stroma 1 and stroma 2 clusters, epithelial clusters, and immune clusters were arranged in layers that are morphologically supported by the anatomical architecture of colonic tissue [Mills, 2019]: (from lumen to serosa) mucosal epithelium; lamina propria (in which immune cells are abundant, and the isolated lymphoid nodules present in this tissue extend into the submucosal layer); submucosal layer, the stromal layer with abundant connective tissue; and lastly muscularis externa, which is represented by the muscle layer.

#### 3.4.2 Pathway analysis of the signature genes identified in the DE analysis

We further conducted pathway analysis using gene ontology [Consortium, 2021] for the signature genes from each cluster identified by SC-MEB. Supplementary Table S9 shows the top four pathways in each cluster. For the regions identified as muscle, muscle contraction was among the top four significant pathways. For the three identified immune clusters, the most significant pathways included humoral immune response and antimicrobial humoral response. For stroma 2 clusters, extracellular structure organization, and external encapsulating structure organization were among the most significant pathways. We also found similar patterns in the heatmap for the normalized expression of signature genes (Fig. 5) between stroma 1 and 2 clusters, among the three immune clusters, and between epithelial clusters 1 and 2. There was high cosine similarity between the two stromal clusters (0.97), as well as among the immune clusters (see Supplementary Table S10). We ultimately compared the mean expression of COVID-19 signature genes [Lee et al., 2020] in the immune and non-immune regions identified by SC-MEB (Fig. 6), and it was clear that COVID-19 signature genes were more highly expressed in the immune regions than the non-immune regions of the colorectal tumor sample.

**Figure 6:**
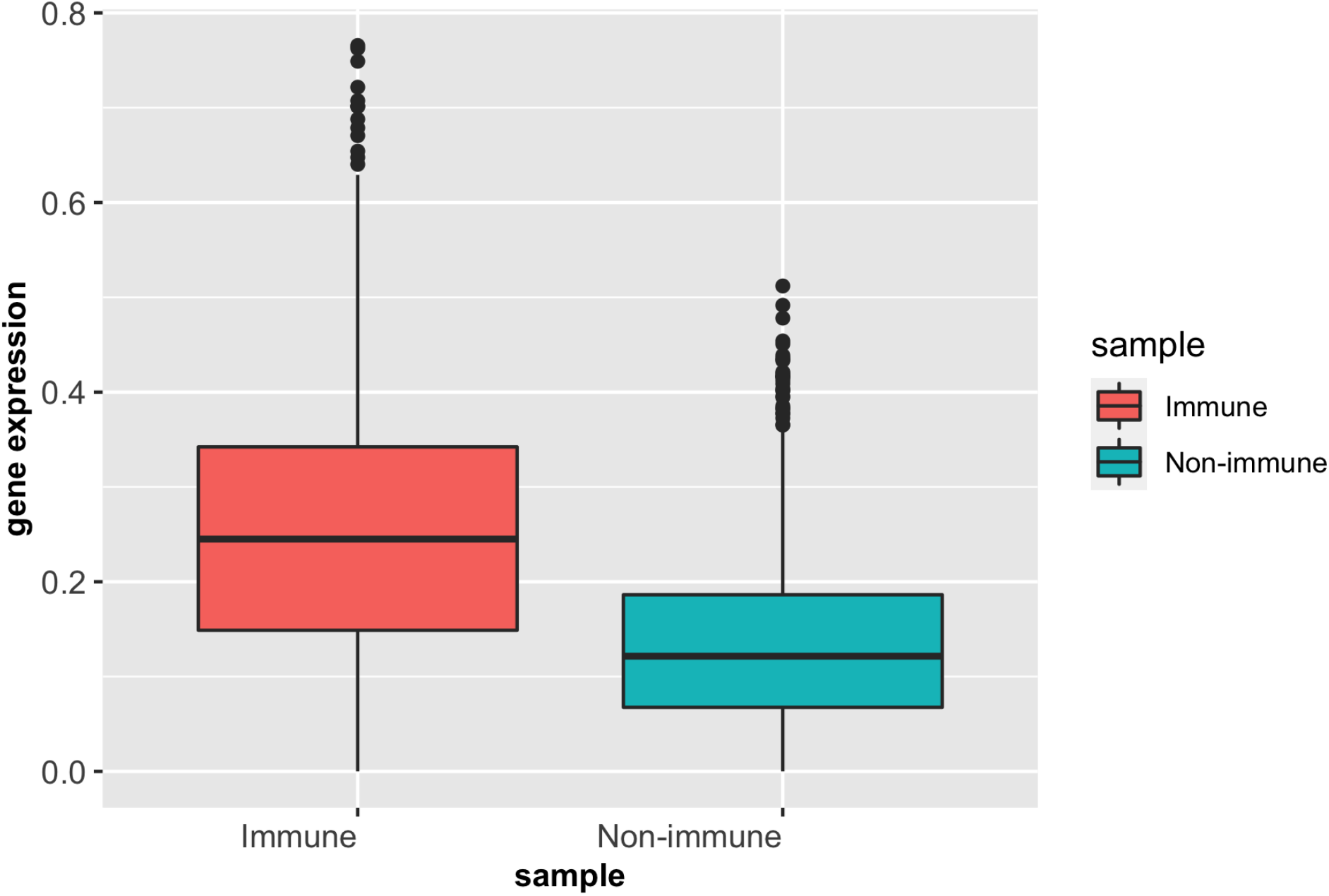
Boxplots of mean expression of COVID-19 signature genes in immune and nonimmune regions.

## 4 Discussion

We propose a new computational tool, SC-MEB, for identifying cell types in spatial transcriptomics (ST) data analysis. Our method builds on a two-level hierarchical probabilistic model that is computationally efficient and can be easily used to analyze the high-resolution data generated by ST technology. Compared with existing approaches, SC-MEB is both more computationally efficient, and is more powerful when there is no prior knowledge regarding the number of clusters is available. We have illustrated the benefits of SC-MEB through extensive simulations, as well as in-depth analysis of four real data sets.

We benchmarked the clustering performance of SC-MEB as well as its computational efficiency using two ST datasets, DLPFC from 10x Genomics Visium and MHPR from MER-FISH, respectively. In the DLPFC dataset, the ARI values for SC-MEB and BayesSpace were comparable, but SC-MEB was 200 times faster at running the analysis than BayesSpace. More importantly, SC-MEB optimized the smoothness parameter *β* and selected the number of clusters in a data-driven manner. SC-MEB could also be applied to perform spatial clustering in other types of ST datasets. Our analysis of the MHPR dataset from MERFISH showed that SC-MEB not only outperformed other methods but was also scalable to larger sample sizes. It took less than five hours to complete the analysis for all cells (*>* 1 million). By applying SC-MEB and other methods, we performed spatial clustering for a colon dataset from a patient with colorectal cancer (CRC) and COVID-19 and further performed differential expression analysis to identify signature genes related to the clustering results. We compared the heatmaps of signature genes identified using SC-MEB and BayesSpace and observed that the clusters identified using SC-MEB were more separable. Using pathway analysis, we identified three immune-related clusters and in a further comparison, we found the mean expression of COVID-19 signature genes was greater in immune than non-immune regions.

There are some caveats associated with SC-MEB that may require further explorations. First, although clustering using low-dimensional features ensures computational efficiency, it is not certain that the features obtained are relevant to class labels that could improve the spatial clustering performance. Thus, an optimal strategy might be to perform joint dimension reduction and spatial clustering for high-dimensional ST datasets. Second, problems with bulk and single-cell RNA-seq remain in the analysis of ST datasets. For example, without removing batch effects from different experiments, findings from differential expression analysis under different conditions could be confounded.

## 5 Competing interests

The authors have no competing interests.

## 6 Author contributions statement

J.L. and J.Y. initiated and designed the study, X.S. and Y.Y. implemented the model and performed simulation studies and benchmarking evaluation, X.S. and J.L. wrote the manuscript, and all authors edited and revised the manuscript.

## 7 Acknowledgments

The authors thank the editor and three anonymous reviewers for their valuable suggestions. This work was supported by grant R-913-200-098-263 from the Duke-NUS Medical School, AcRF Tier 2 (MOE2018-T2-1-046 and MOE2018-T2-2-006) from the Ministry of Education, Singapore, grants (NSF: # 1636933 and # 1920920) from the National Science Foundation of China.

## Notes

https://github.com/Shufeyangyi2015310117/SC.MEB

### Competing Interest Statement

The authors have declared no competing interest.

