## supplementary for "SC-MEB: spatial clustering with hidden Markov random field using empirical Bayes"

#### 1 SC-MEB Method

##### 1.1 the Hidden Markov random field model

Let $\mathcal{S}$ be the set of all spots with a neighborhood system defined on it. The cell types in each spot can be modeled by a hidden random field

$$\{x = (x_1, \ldots, x_N) \mid x_i \in \mathcal{L}, i \in \mathcal{S}\},$$

where $N$ is the number of spots, and $\mathcal{L} = \{1, \ldots, K\}$ is the set of all possible cell types. The $p$-dimensional representation of the gene expression is modeled by an observed random field

$$\{y = (y_1, \ldots, y_N) \mid y_i \in \mathbb{R}^p, i \in \mathcal{S}\}.$$

For spot $i$, its neighbors we considered are the spots surrounding it, and is denoted by $\mathcal{N}_i$.

The representation of the gene expression $y$ is assumed to be a Gaussian MRF model, with one prefixed parameters, the number of cell types $K$. Specifically, the observed random field $y$ are assumed to be mutually independent given the hidden random field $x$, and normally distributed with:

$$p(y \mid x, \theta) = \prod_{i \in \mathcal{S}} \mathcal{N}(y_i | x_i = k, \mu_k, \Sigma_k),$$

where $\theta = \{\mu_k, \Sigma_k : k = 1, \ldots, K\}$, $\mu_k$ and $\Sigma_k$ denote the mean and Covariance matrix for cell type $k$. The hidden random field $x$ are assumed to be the Potts model:

$$\begin{aligned} P(x) &= \frac{1}{Z_\beta} \exp\{-U(x)\}, \\ U(x) &= \sum_{i, i' \in \mathcal{N}_i} \beta[1 - \delta(x_i, x_i')], \end{aligned} \tag{1}$$

where $\delta$ is the delta function, and $Z_\beta$ is a normalization constant that lacks a closed form.

With prefixed $K$, the question of interest is to recover the unknown cell types $x$, interpreted as a clustering into a finite number $K$ of types. This clustering usually is conducted based on the posterior distribution of $x$, and requires the estimation of parameter $\phi = (\theta, \beta)$.

#### 1.2 ICM-EM algorithm

Estimation of the parameter is done using an iterative conditional mode based expectation-maximization (ICM-EM) algorithm [Cuadra et al., 2005]. We assume $K$ is known at present, and will discuss their selection in next section.

In ICM step, the estimate of $x$ is obtained by maximizing its posterior with respect to $x_i$ coordinately:

$$P(x \mid y) = P(x_i, x_{\mathcal{S}-\{i\}} \mid y) = P(x_i \mid y, x_{\mathcal{S}-\{i\}}) P(x_{\mathcal{S}-\{i\}} \mid y),$$

where $i = 1, \ldots, n$, until converge [Besag, 1986]. Given initial values of $x$, $\theta$, and observed $y$, we have the update equation:

$$\hat{x}_i = \min_{x_i} V(\hat{x}_1, \ldots, x_i, \ldots, \hat{x}_n), \tag{2}$$

where

$$\begin{aligned} V(x) = &\Big\{ \frac{1}{2}(y_i - \mu_{x_i})^\top \Sigma_{x_i}^{-1} (y_i - \mu_{x_i}) + \frac{1}{2} \log |\Sigma_{x_i}| \\ &+ \beta \sum_{i' \in \mathcal{N}_i} [1 - \delta(x_i, x_{i'})] \Big\}. \end{aligned}$$

In EM step, instead of the original complete likelihood which is difficult to perform the E-step, the following pseudo-likelihood is used:

$$\begin{aligned}\tilde{p}(y, x; \phi) &= p(y|x; \phi)\tilde{p}(x; \phi)\\ &= \prod_i p(y_i \mid x_i; \phi) \prod p(x_i|x_{\mathcal{N}_i}; \phi)\\ &= \prod_i [p(y_i \mid x_i; \phi)p(x_i|x_{\mathcal{N}_i}; \phi)]\\ &= \prod_i p(y_i, x_i \mid x_{\mathcal{N}_i}; \phi).\end{aligned}$$

For any $q_i(x_i)$, we have

$$\begin{aligned}\log \tilde{p}(y; \phi) &= \sum_i \log \sum_k p(y_i, x_i = k \mid x_{\mathcal{N}_i}; \phi)\\ &= \sum_i \log \sum_k q_i(x_i = k) \frac{p(y_i, x_i = k \mid x_{\mathcal{N}_i}; \phi)}{q_i(x_i = k)}\\ &= \sum_i \log \mathbf{E}_{x_i} \left[\frac{p(y_i, x_i \mid x_{\mathcal{N}_i}; \phi)}{q_i(x_i)}\right]\\ &\geq \sum_i \mathbf{E}_{x_i} \left[\log \frac{p(y_i, x_i \mid x_{\mathcal{N}_i}; \phi)}{q_i(x_i)}\right] := \text{ELBO}(\phi).\end{aligned}$$

The above inequality is an equality if and only if

$$q_i(x_i) = \frac{p(y_i, x_i \mid x_{\mathcal{N}_i}; \phi)}{\sum_{x_i} p(y_i, x_i \mid x_{\mathcal{N}_i}; \phi)} = p(x_i \mid y_i, x_{\mathcal{N}_i}).$$

Then we have

$$\begin{aligned}\text{ELBO}(\phi) =& \sum_i \sum_k \gamma_{ik} \log p(y_i, x_i = k \mid x_{\mathcal{N}_i}; \phi) - \sum_i \sum_k \gamma_{ik} \log \gamma_{ik}\\ =& \sum_i \sum_k \gamma_{ik} [\log p(y_i \mid x_i = k; \phi) + \log p(x_i = k \mid x_{\mathcal{N}_i}; \phi)]\\ & - \sum_i \sum_k \gamma_{ik} \log \gamma_{ik},\\ :=& Q(\phi) + \text{Const},\end{aligned}$$

where $\gamma_{ik}$ is called responsibility that component $k$ takes for explaining the observation $y_i$, and is defined as follows:

$$\gamma_{ik} = \frac{P(y_i \mid x_i = k)P(x_i = k \mid X_{\mathcal{N}_i} = \hat{x}_{\mathcal{N}_i})}{\sum_{k'} P(y_i \mid x_i = k')P(X_i = k' \mid X_{\mathcal{N}_i} = \hat{x}_{\mathcal{N}_i})}. \tag{3}$$

By taking partial derivatives of $Q(\phi)$ with respect to the parameters and setting them to zero, we obtain the update equations for the maximization step:

$$\mu_k = \frac{1}{N_k} \sum_{i=1}^{n} \gamma_{ik} y_i, \tag{4}$$

$$\Sigma_k = \frac{1}{N_k} \sum_{i=1}^{n} \gamma_{ik} (y_i - \mu_k)(y_i - \mu_k)^\top, \tag{5}$$

$$\tag{6}$$

where $N_k = \sum_{i=1}^{n} \gamma_{ik}$. Since there is no closed-form solution for $\hat{\beta}$, we optimize the smoothness parameter $\beta$ via a grid search strategy:

$$\beta = \arg \max_{l \in \{1,\ldots,R\}} Q(\theta, \beta_l), \tag{7}$$

where the sequence $(\beta_1, \ldots, \beta_R)$ is a vector of 20 evenly spaced points in the interval $[0, 4]$.

The ICM-EM algorithm iterates the ICM step and maximization steps until convergence. Implementations are summarized in Algorithm 1 for clarity.

**Algorithm 1:** ICM-EM with prefixed $K$

```
Initialize x, φ = {μ_k, Σ_k : k = 1, ..., K, β}.
repeat
    ICM-step:
    while the change of V(x) is larger than a threshold do
        for i = 1, ..., n do
            x̂_i = min_{x_i} V(x̂_1, ..., x_i, ..., x̂_n), ;
        end
    end
    E-step:
    for i = 1, ..., n, k = 1, ..., K do
        γ_ik = P(y_i|x_i=k)P(x_i=k|X_{N_i}=x̂_{N_i}) / Σ_{k'} P(y_i|x_i=k')P(X_i=k'|X_{N_i}=x̂_{N_i}) ;
    end
    M-step:
    for k = 1, ..., K do
        N_k = Σ_{i=1}^n γ_ik;
        μ_k = (1/N_k) Σ_{i=1}^n γ_ik y_i;
        Σ_k = (1/N_k) Σ_{i=1}^n γ_ik (y_i − μ_k)(y_i − μ_k)^⊤;
        β = arg max_{l∈{1,...,R}} Q(θ, β_l);
    end
until the change of log p̃(y; φ) is smaller than a threshold;
return x̂ and φ.
```

Where the listing's math is: $\hat{x}_i = \min_{x_i} V(\hat{x}_1, \ldots, x_i, \ldots, \hat{x}_n)$; $\gamma_{ik} = \frac{P(y_i|x_i=k)P(x_i=k|X_{\mathcal{N}_i}=\hat{x}_{\mathcal{N}_i})}{\sum_{k'} P(y_i|x_i=k')P(X_i=k'|X_{\mathcal{N}_i}=\hat{x}_{\mathcal{N}_i})}$; $N_k = \sum_{i=1}^{n} \gamma_{ik}$; $\mu_k = \frac{1}{N_k}\sum_{i=1}^{n} \gamma_{ik} y_i$; $\Sigma_k = \frac{1}{N_k}\sum_{i=1}^{n} \gamma_{ik}(y_i - \mu_k)(y_i - \mu_k)^\top$; $\beta = \arg\max_{l\in\{1,\ldots,R\}} Q(\theta, \beta_l)$; $\log \tilde{p}(y; \phi)$; $\hat{x}$ and $\phi$.

#### 1.3 Spatial clustering model selection with modified BIC (MBIC)

Since the number of cell types is not known in advance, we use the modified BIC (MBIC) [Wang et al., 2007] to choose it.

Let $M_r$ denote a MRF model with prefixed $K_r$, where $K_r = K_{\min}$ to $K_r = K_{\max}$. Denote $\phi_r$ the parameters in $M_r$. The modified BIC is defined as follows [Wang et al., 2007]:

$$\mathrm{MBIC}(\hat{\phi}_r) = 2 \log p(y \mid \hat{\phi}_r) - C_n d_r \log n \quad (8)$$

where $d_r$ is the number of free parameters in $M_r$, and equals to $K \times p + K \times \frac{p(p+1)}{2}$ in a K-labels MRF model. The positive constant $C_n = c \log(\log(n + p))$ [Ma and Huang, 2017], where $c$ is specified by users, and usually can be set 1, 5 or 10. Note that when the $C_n$ equals to 1, the MBIC reduces to BIC. Here the logarithm of the observed likelihood $p(y \mid \hat{\phi}_r)$ is approximated with the $\log \tilde{p}(y; \hat{\phi}_r)$.

### 2 Simulation Results

#### 2.1 Additional results for Example I

In Example I, lables of spots are radomly generated. We generated 4900 spatial spots defined on a $70 \times 70$ square lattices, and simulated the cluster label for each spot from the $K$-states Potts model with $\beta \in [1, 1.3]$ using the R package *GiRaF*. The number of neighbors is set to 4 and the number of true clusters $K$ is set to be 3, 5 or 7. At last, two distributions are considered for PC representations of gene expression: a Gaussian mixture model, and a Student-t mixture model. The number of PCs was set to be either 10, or 15. The component mean $\mu_k$ and the component covariance matrix $\Sigma_k$ are described in Supplementary Table S1-S4.

**Figure S1.** All component densities have different covariances, and the number of PCs was set to be 10.

**Figure S2.** All component densities have different covariances, and the number of PCs was set to be 15.

**Figure S3.** All components have a shared covariance, and the number of PCs was set to be 10.

**Figure S4.** All components have a shared covariance, and the number of PCs was set to be 15.

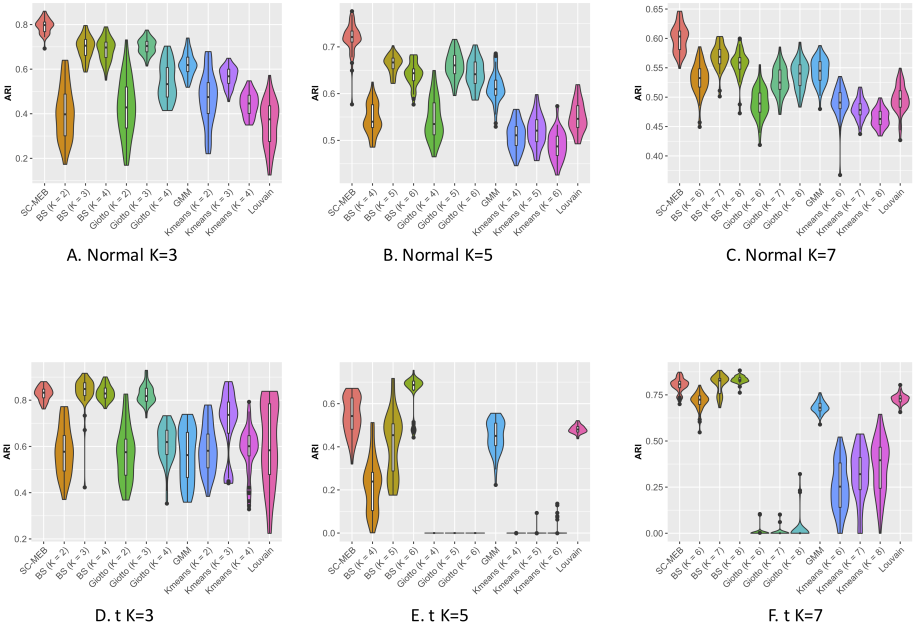


Figure S1: Additional results for Example I. All component densities have different covariances, and the number of PCs was set to be 10. **A-C.** PCs are sampled from a Gaussian mixture model, and the number of true clusters $K$ is set to be be 3, 5 or 7. **D-F.** PCs are sampled from a Student-$t$ mixture model, and the number of true clusters $K$ is set to be be 3, 5 or 7.

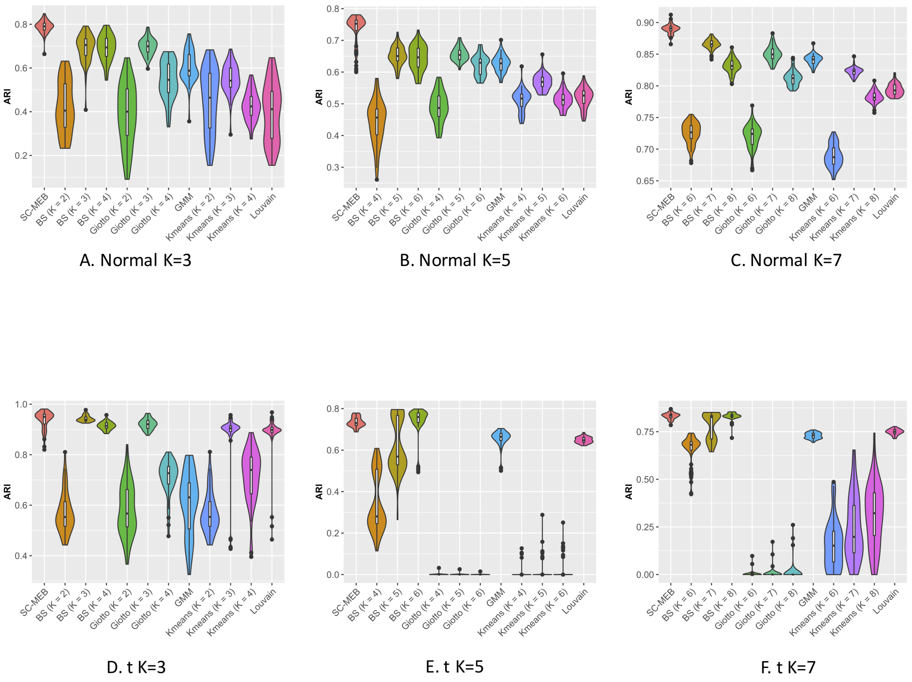


Figure S2: Additional results for Example I. All component densities have different covariances, and the number of PCs was set to be 15. **A-C.** PCs are sampled from a Gaussian mixture model, and the number of true clusters $K$ is set to be be 3, 5 or 7. **D-F.** PCs are sampled from a Student-$t$ mixture model, and the number of true clusters $K$ is set to be be 3, 5 or 7.

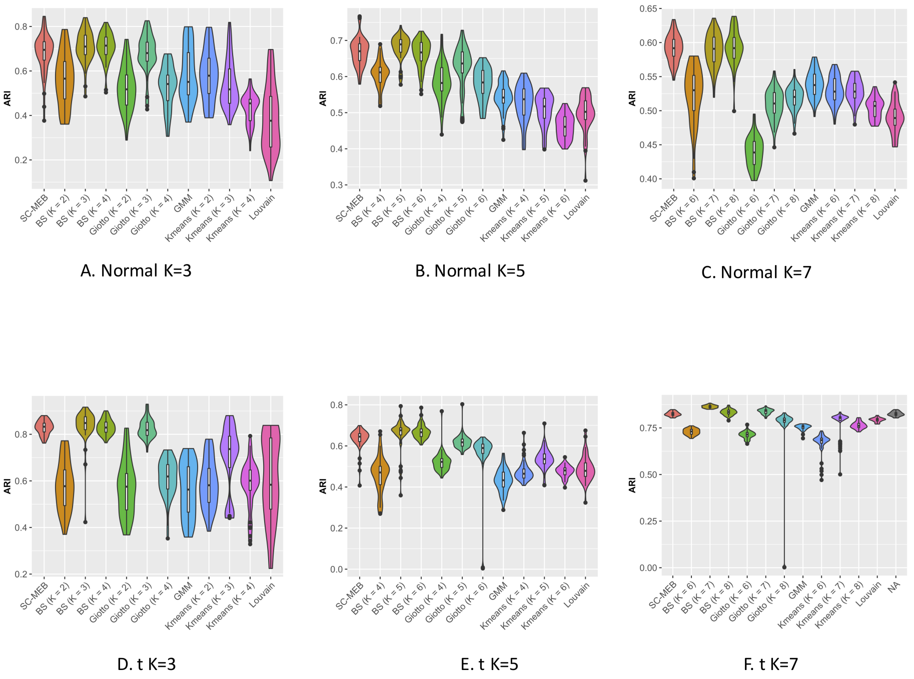


Figure S3: Additional results for Example I. All components have a shared covariance, and the number of PCs was set to be 10. **A-C.** PCs are sampled from a Gaussian mixture model, and the number of true clusters $K$ is set to be be 3, 5 or 7. **D-F.** PCs are sampled from a Student-$t$ mixture model, and the number of true clusters $K$ is set to be be 3, 5 or 7.

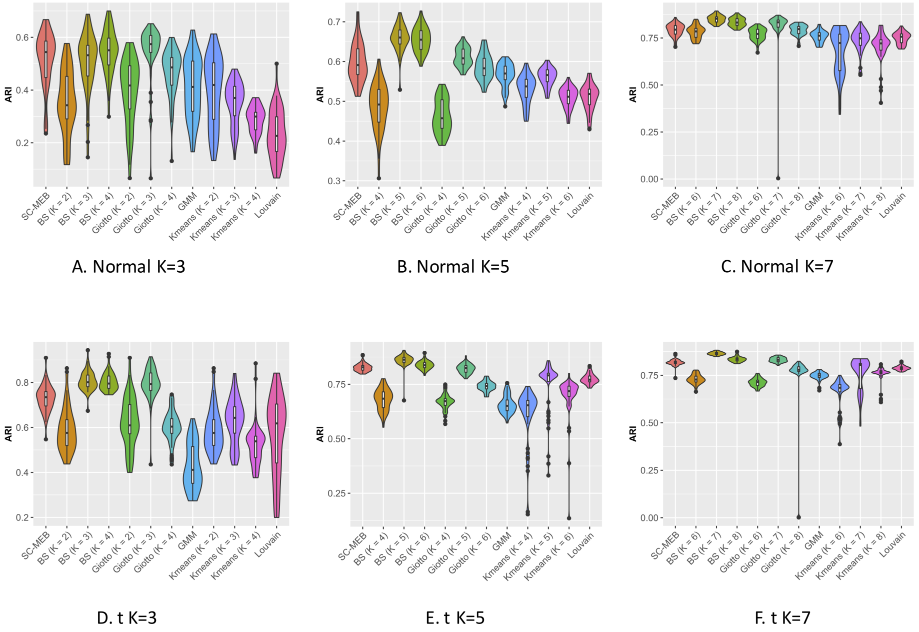


Figure S4: Additional results for Example I. All components have a shared covariance, and the number of PCs was set to be 15. **A-C.** PCs are sampled from a Gaussian mixture model, and the number of true clusters $K$ is set to be be 3, 5 or 7. **D-F.** PCs are sampled from a Student-$t$ mixture model, and the number of true clusters $K$ is set to be be 3, 5 or 7.

#### 2.2 Additional results for Example II

In Example II, lables of spots are obtained from our real data. Specifically, we obtained labels of 2,988 spots in the CRC data inferred from the corresponding real data with SC-MEB ($K = 8$). PCs were sampled in the same way as Example I. The component mean $\mu_k$ and the component covariance matrix $\Sigma_k$ are described in Supplementary Table S5-S6.

**Figure S5.** The number of PCs was set to be 10.

**Figure S6.** The number of PCs was set to be 15.

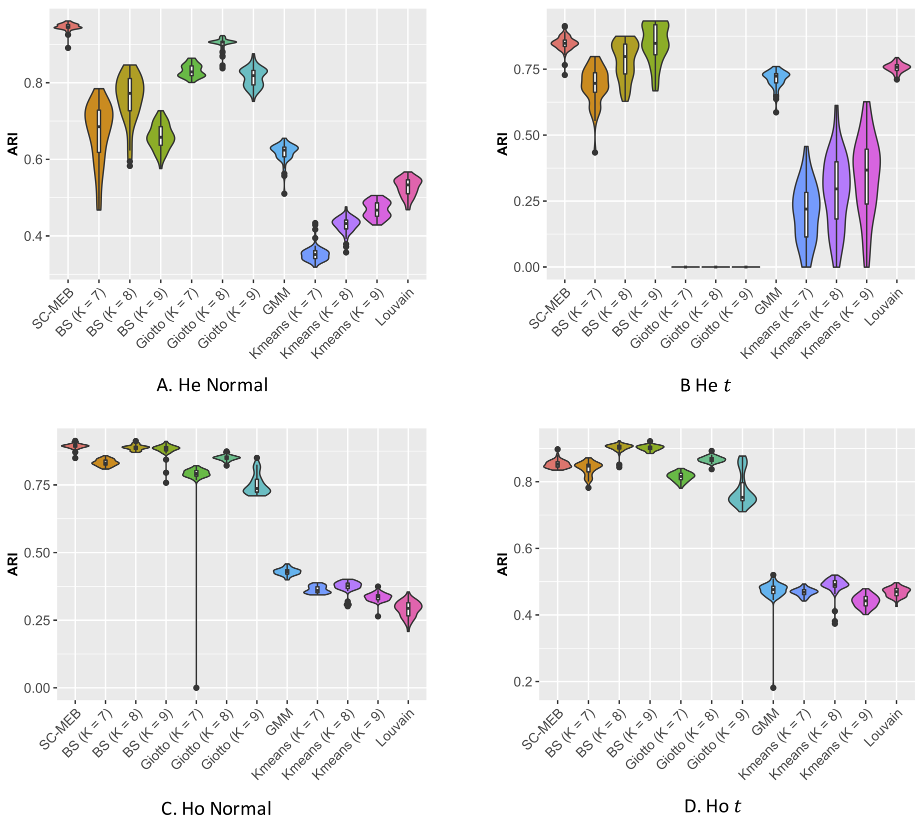


Figure S5: Additional results for Example II. The number of PCs was set to be ten. "He" means all component densities have different covariance matrices, and "Ho" means all component densities have a shared covariance matrix **A-B.** PCs are sampled from a Gaussian mixture model for A and Student's-$t$ mixture model for B; all component densities have different covariances. **C-D.** PCs are sampled from a Gaussian mixture model for C and Student's-$t$ mixture model for D; all components have a shared covariance.

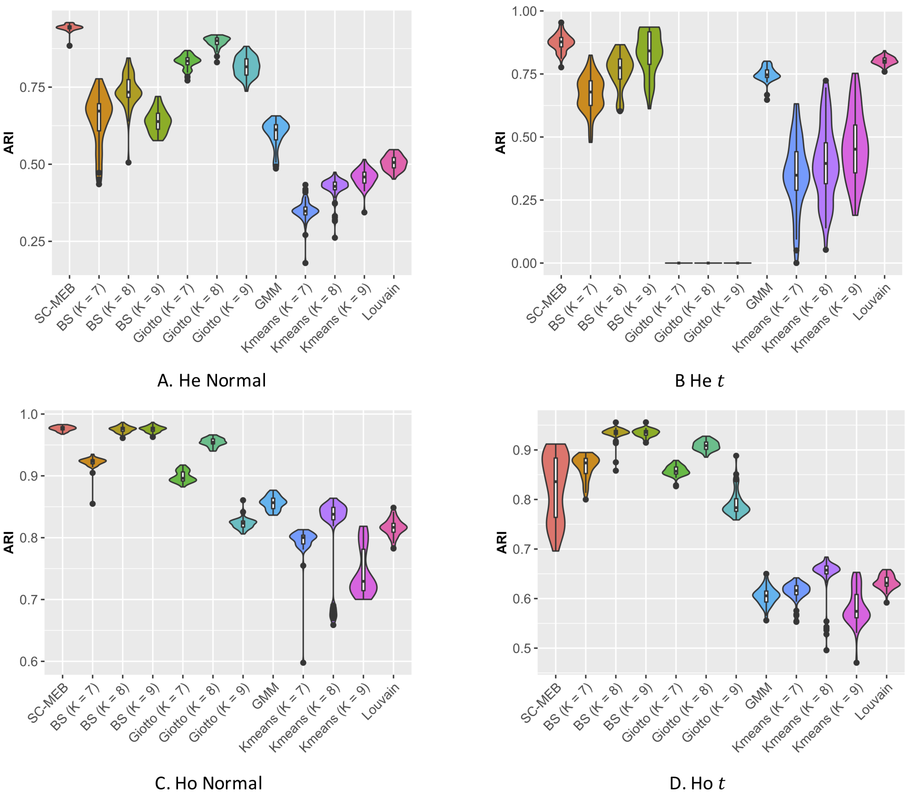


Figure S6: Additional results for Example II. The number of PCs was set to be 15. “He” means all component densities have different covariance matrices, and “Ho” means all component densities have a shared covariance matrix. **A-B.** PCs are sampled from a Gaussian mixture model for A and Student-$t$ mixture model for B; all component densities have different covariances. **C-D.** PCs are sampled from a Gaussian mixture model for C and Student’s-$t$ mixture model for D; all components have a shared covariance.

#### 2.3 The results for scenario when the degrees of freedom (df) of t distribution is small

In this scenario, the lables of spots were obtained in the same way as Example II, a total of 2988 spots and 8 clusters. However, the PCs were sampled from the t distributions with small degrees of freedom. The mean $\mu_k$ and the degrees of freedom $(df_k)$ are described in Supplementary Table S11.

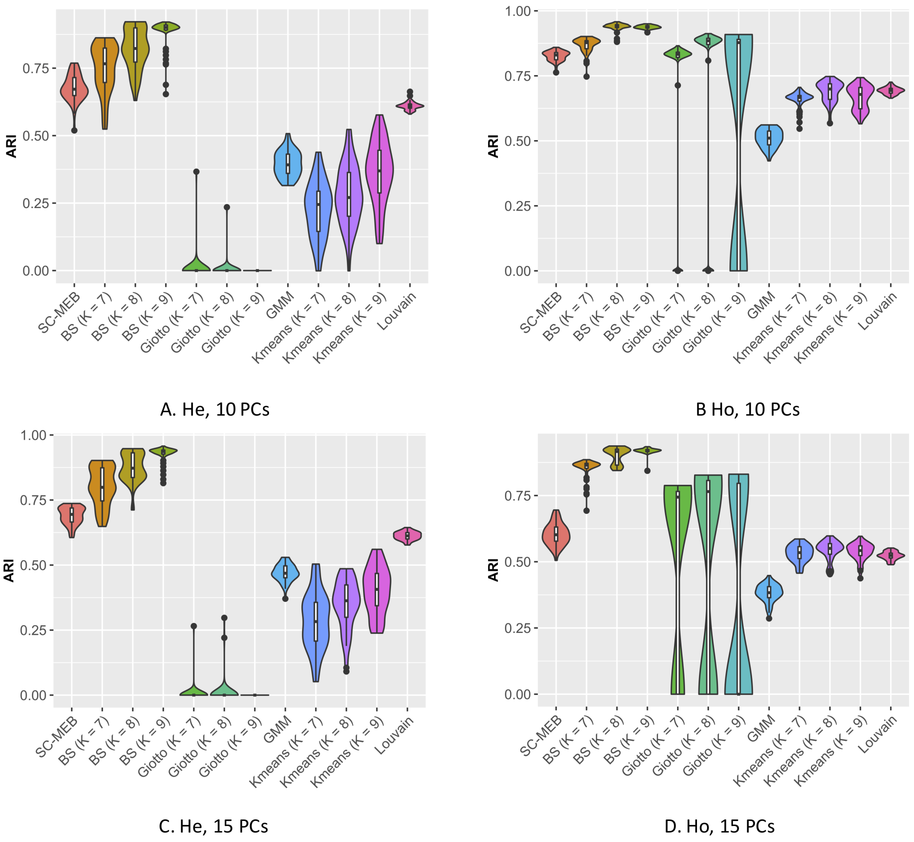


Figure S7: The results for scenario when the degrees of freedom $(df_k)$ of t distribution is small. "He" means all component densities have different covariance matrices, and "Ho" means all component densities have a shared covariance matrix. **A-B.** The number of PCs was set to be 10. **C-D.** The number of PCs was set to be 15.

#### 3 The annotation results for muscle region

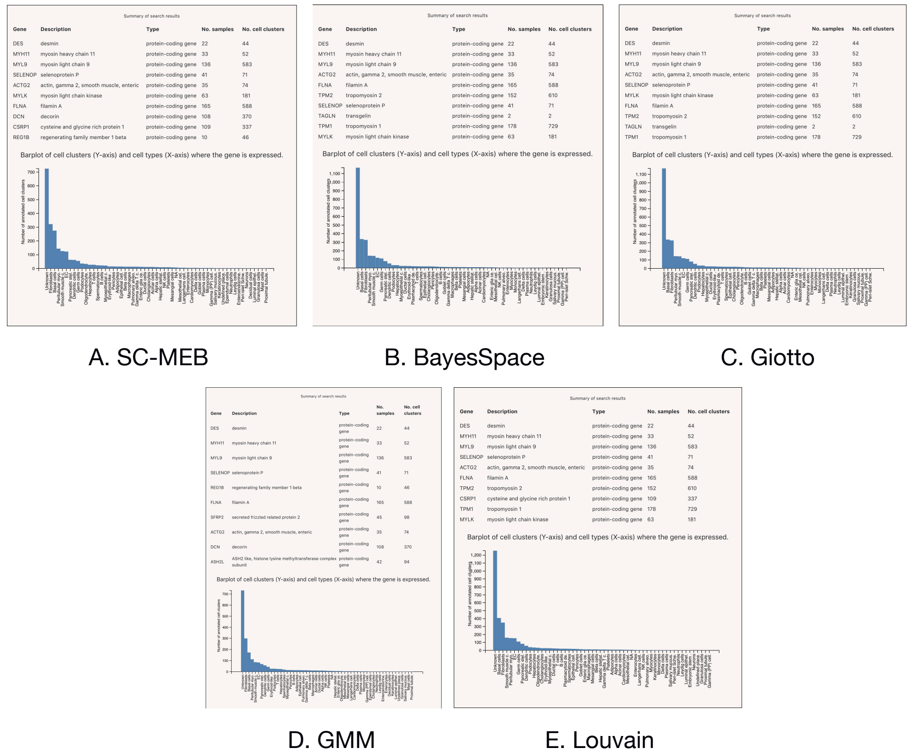


Figure S8: The annotation results for muscle region identified by SC-MEB (A), BayesSpace (B), Giotto (C), GMM (D) and Louvain (E) based on PanglaoDB.
